# AAV11 permits efficient retrograde targeting of projection neurons

**DOI:** 10.1101/2022.01.13.476170

**Authors:** Zengpeng Han, Nengsong Luo, Jiaxin Kou, Lei Li, Wenyu Ma, Siqi Peng, Zihong Xu, Wen Zhang, Yuxiang Qiu, Yang Wu, Jie Wang, Chaohui Ye, Kunzhang Lin, Fuqiang Xu

**Affiliations:** Key Laboratory of Magnetic Resonance in Biological Systems, State Key Laboratory of Magnetic Resonance and Atomic and Molecular Physics, National Center for Magnetic Resonance in Wuhan, Wuhan Institute of Physics and Mathematics, Innovation Academy for Precision Measurement Science and Technology, Chinese Academy of Sciences-Wuhan National Laboratory for Optoelectronics, Wuhan, 430071, P.R. China; University of Chinese Academy of Sciences, Beijing, 100049, P.R. China; The Brain Cognition and Brain Disease Institute (BCBDI), Shenzhen Key Laboratory of Viral Vectors for Biomedicine, Shenzhen Institute of Advanced Technology, Chinese Academy of Sciences; Shenzhen-Hong Kong Institute of Brain Science-Shenzhen Fundamental Research Institutions, NMPA Key Laboratory for Research and Evaluation of Viral Vector Technology in Cell and Gene Therapy Medicinal Products, Shenzhen, Key Laboratory of Quality Control Technology for Virus-Based Therapeutics, Guangdong Provincial Medical Products Administration, Shenzhen, 518055, P.R. China; Wuhan National Laboratory for Optoelectronics, Huazhong University of Science and Technology, Wuhan 430074, P.R. China; Department of Pathophysiology, Key Lab of Neurological Disorder of Education Ministry, School of Basic Medicine, Tongji Medical College, Huazhong University of Science and Technology, Wuhan, P. R. China; College of Life Sciences, Wuhan University, Wuhan, P.R. China; Shenzhen-Hong Kong Institute of Brain Science-Shenzhen Fundamental Research Institutions, Shenzhen, 518055, P.R. China; Center for Excellence in Brain Science and Intelligence Technology, Chinese Academy of Sciences, Shanghai, 200031, P.R. China

**Keywords:** Adeno-associated virus 11, Retrograde, Natural serotype capsid, Projection Neurons

## Abstract

Viral tracers that permit efficient retrograde targeting of projection neurons are powerful vehicles for structural and functional dissections of the neural circuit and for the treatment of brain diseases. Recombinant adeno-associated viruses (rAAVs) are the most potential candidates because they are low-toxic with high-level transgene expression and minimal host immune responses. Currently, some rAAVs based on capsid engineering for retrograde tracing have been widely used in the analysis and manipulation of neural circuits, but suffer from brain area selectivity and inefficient retrograde transduction in certain neural connections. Here, we discovered that the recombinant adeno-associated virus 11 (rAAV11) exhibits potent retrograde labeling of projection neurons with enhanced efficiency to rAAV2-retro in some neural connections. Combined with calcium recording technology, rAAV11 can be used to monitor neuronal activities by expressing Cre recombinase or calcium-sensitive functional probe. In addition, we further showed the suitability of rAAV11 for astrocyte targeting. These properties make rAAV11 a promising tool for the mapping and manipulation of neural circuits and gene therapy of some neurological and neurodegenerative disorders.

**Highlights:**

1. Naturally occurring AAV serotype capsid exhibits robust retrograde functionality
2. Improved distribution properties and retrograde transport efficiency
3. Can express Cre recombinase or calcium-sensitive functional probe for neural circuits monitoring
4. Can specifically target astrocytes

## Introduction

Several major mental diseases caused by structural variation or loss of neural circuits, such as Parkinson’s disease, Alzheimer’s disease, schizophrenia, autism, etc., have brought a heavy burden to the family and society. At present, there is still a lack of effective prevention and cure methods, and the main reason is that the mechanism of disease-related complex circuit changes is unclear. Analyzing the structure and function of brain neural network is fundamental to revealing the working principle of brain and the mechanism of brain diseases. However, mapping and manipulating functional network connection has become an important challenge due to the lacking of techniques available for targeting projection neurons. While classic neuroanatomical retrograde tracers have been widely used to analysis the architectural connectivity between different brain regions, they cannot deliver genetic cargos for neural activity manipulation and clinical application [1].

Recombinant viral vectors have been developed as valuable tools to address the limitations of traditional retrograde tracers, because they can not only express fluorescent proteins for connection network visualization, but also carry functional probes (e.g., indicators or effectors of neural activity) for the monitoring or manipulation of neural circuits. Many naturally evolved and engineered viruses can transduce projection neuron populations by axon terminal uptake [2, 3], including canine adenovirus-2 (CAV-2), rabies virus (RV), herpes simplex virus (HSV), and lentivirus (LV) among others. Of these, CAV-2 can effectively retrograde label long-range projection neurons [4, 5], but allows only moderate levels of transgene expression, presents some toxicity [6, 7], and has difficulties in large-scale preparation for clinic applications [8]. While RV and HSV display robust efficiency of retrograde labeling with rapid gene expression, they are not suitable for functional research and gene therapy due to high cytotoxicity. [9–14]. Lentiviruses pseudotyped with modified RVG present the improved retrograde transduction ability in rodents and non-human primates [15, 16], but have a risk of tumorigenesis due to random integration into the host genome [17]. Thus, developing non-toxic, readily manufactured viral vectors that mediate flexible packaging of different transgenes, retrograde transduction, and high-level gene expression remains an urgent need.

In this case, the use of capsid engineering adeno-associated virus (AAV) for retrograde labeling becomes a good option because of its safety, non-pathogenic nature, and ability to infect dividing and quiescent cells (e.g., neurons) *in vivo* [18, 19]. Many engineered AAV capsids for retrograde tracing have been reported, such as AAV2-retro [20], AAV9-retro [21], AAV2-MNM004 [22], AAV9-SLR [23], etc. Among them, AAV2-retro was developed by the insertion of a fragment (LADQDYTKTA) from 10-mer peptide libraries between N587 and R588 of the wild-type AAV2 VP1 capsid gene, displaying the most robust retrograde transport, has been extensively used to express fluorescent probes to analyze the structural connections of neural networks and to express functional probes to monitor and manipulate neuronal activities [20]. However, AAV2-retro suffers from brain area selectivity and inefficient retrograde transduction in certain neural connections [24]. And because the retrograde characteristics of these AAV carriers are given through technical modification, it may not be easy to have further improvements.

In addition to the capsid engineering methods such as directed evolution or rational design, a more fundamental way to find a better AAV is to identify new natural serotypes or new characteristics of natural serotypes. The results of this method can be used as a basis for further engineering work, thereby increasing the underlying optimization potential. There are many natural serotypes of AAV, which belong to different pedigrees and have their unique infection characteristics [25–27]. In neuroscience research, the commonly used AAV serotypes are 1, 2, 5, 6, 8, and 9, but low propensity of retrograde transduction hinders their applications in the research of projection network or disease treatment. Here, we developed the recombinant adeno-associated virus serotype 11 (rAAV11), and found that it exhibits potent retrograde labeling of projection neurons with enhanced efficiency to rAAV2-retro in some neural connections. Combined with calcium recording technology, rAAV11 can be used to monitor neuronal activities by expressing Cre recombinase or calcium-sensitive functional probe. In addition, we further showed the suitability of rAAV11 for astrocyte targeting. These properties make rAAV11 a promising tool for the mapping and manipulation of neural circuits and gene therapy of some neurological and neurodegenerative disorders. As a basis vector, it has considerable potential for further engineering optimization.

## Results

### Obtain rAAV11 virus vector based on HEK293T production system

We have previously established a single baculovirus-insect cell-based large-scale production system (OneBac system) and verified the preparation of rAAV1-13 serotypes [28, 29]. Although it is conducive to large-scale production, putting all the genetic elements in the common bacmid is not convenient for switching the carried foreign genes. To facilitate laboratory-level verification, we established the AAV packaging method based on the three plasmids. The pAAV2/11 plasmid was obtained by inserting AAV11 *Cap* sequence into the pAAV2/1 backbone to substitute the AAV1 *Cap* gene and its efficiency in virus packaging was evaluated in HEK-293 cells. We found that pAAV2/11 plasmid could be used to package high-titer rAAV2/11 with efficiency comparable to rAAV2/9.

### rAAV11 mediates efficient neuron transduction with axon terminal absorption

The dorsal striatum, called the caudate-putamen (CPu), includes the caudate and putamen nucleus, which is innervated by dopaminergic neurons from the substantia nigra (SN) pars compacta [30]. It’s one of the structures that compose the basal nuclei. Through various pathways, it receives dopaminergic inputs from the ventral tegmental area (VTA) and the SNr and glutamatergic inputs from several regions, including the cortex, hippocampus, amygdala, and thalamus [31–35]. A primary function of the caudate-putamen is to regulate movements at various stages and influence different types of learning. The caudate-putamen also plays a role in degenerative neurological disorders, such as Parkinson’s disease [36].

To evaluate the infectious effect of rAAV11, rAAV11-EF1α-EGFP viruses were delivered to CPu via stereotactic injection [3 × 10^9^ vector genomes (VG) in total] in adult mice and the fluorescence distribution was assessed (Fig.1 A, B). Fig.1 B shows the fluorescence distribution of EGFP expression at the injection site, with a pretty broad spread range. Fig.1 C shows the spread of the virus towards the anterior side, Fig.1 D shows the spread of the virus to the posterior side, clear neuron morphology can be labeled in several upstream brain areas, such as somatomotor areas (MO), anterior olfactory nucleus (AON), primary somatosensory area (SSp), and mediodorsal nucleus of thalamus (MD), among others. These results suggest that rAAV11 has excellent retrograde labeling capability and can be transported to the cellular body of upstream neurons by axon terminal uptake.

**Fig.1.**
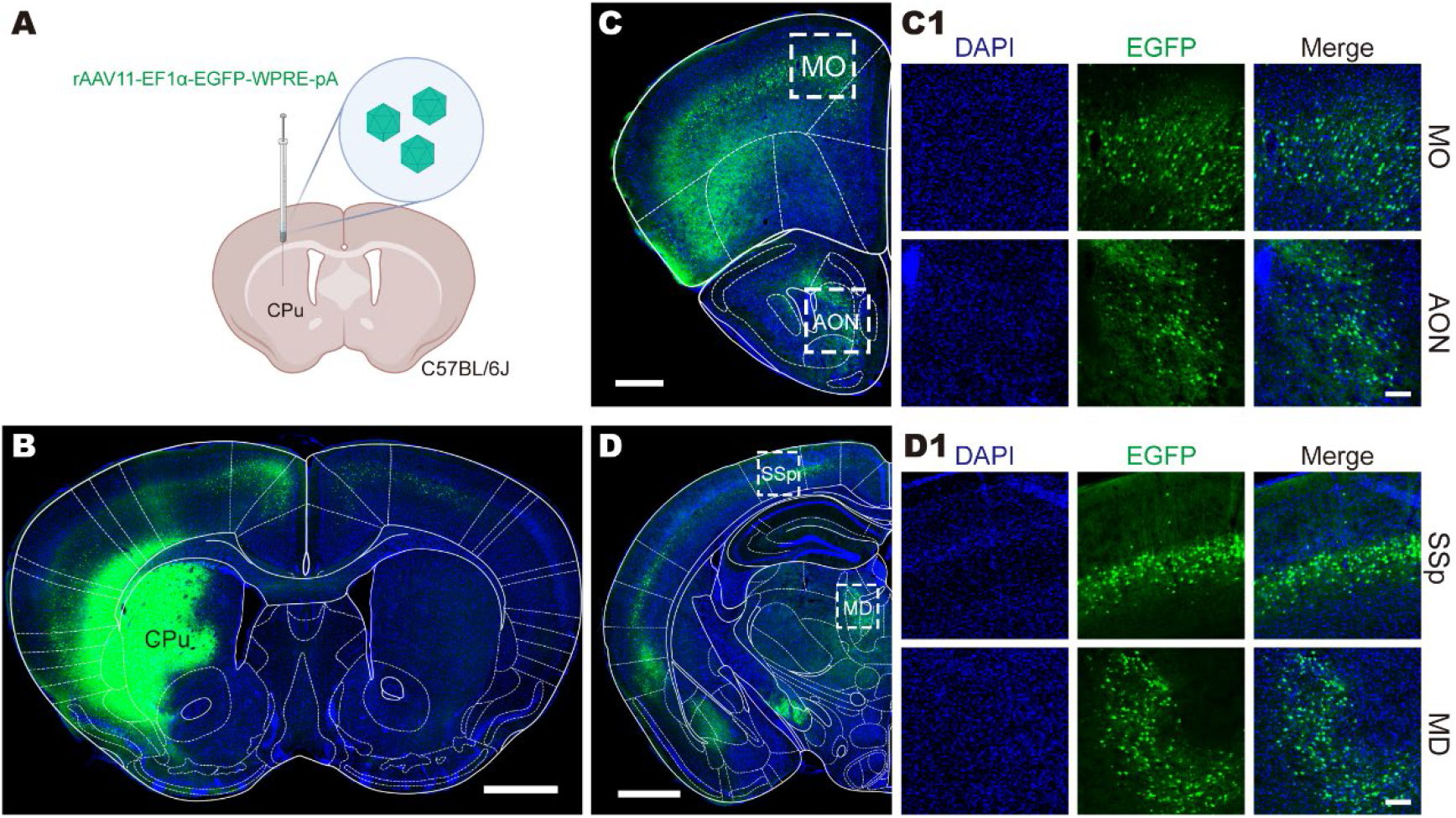
Natural serotype AAV11 capsid for efficient retrograde access to projection neurons. (A) Schematic diagram of virus injection, rAAV11-EF1α-EGFP-WPRE-pA (3 × 10^9^ VG) were injected into caudate-putamen (CPu) area in C57BL/6J mice. (B) The fluorescence distribution of EGFP expression in injection site CPu. Scale bar = 1 mm. (C) The spread of the virus to the anterior side. Scale bar = 500 μm. (C1) Neurons were tagged in somatomor areas (MO) and anterior olfactory nucleus (AON). Scale bar = 100 μm. (D) The spread of the virus to the posterior side, Scale bar = 1 mm. (D1) Neurons were tagged in somatosensory area (SSp) and mediodorsal nucleus of thalamus (MD). Scale bar = 100 μm.

### rAAV11 does not exhibit anterograde transsynaptic spread property

Some AAV serotypes (such as AAV1 and AAV9) exhibit the features of anterograde transsynaptic propagation [37], which can also cause an extensive spread range. To test whether rAAV11 can mediate anterograde transsynaptic spread, rAAV11-hSyn-Cre was injected into primary visual cortex (V1) of Ai14 (CAG promoter-driven and Cre-dependent expression of tdTomato reporter) mice [38] (2 × 10^9^ VG in total) (Fig.2 A, B). Following a 3 week post-injection, a number of tdTomato fluorescence signals were detected at the injection site, but no tdTomato-expressing cell bodies were observed in the downstream regions known to be directly projected by V1 (Fig.2 B, B1, C, C1), including the superior colliculus (SC) and caudate-putamen (CPu), which had been proved not to project to V1 [39, 40], indicating that the spread of rAAV11 is not caused by the anterograde transsynaptic spread.

**Fig.2.**
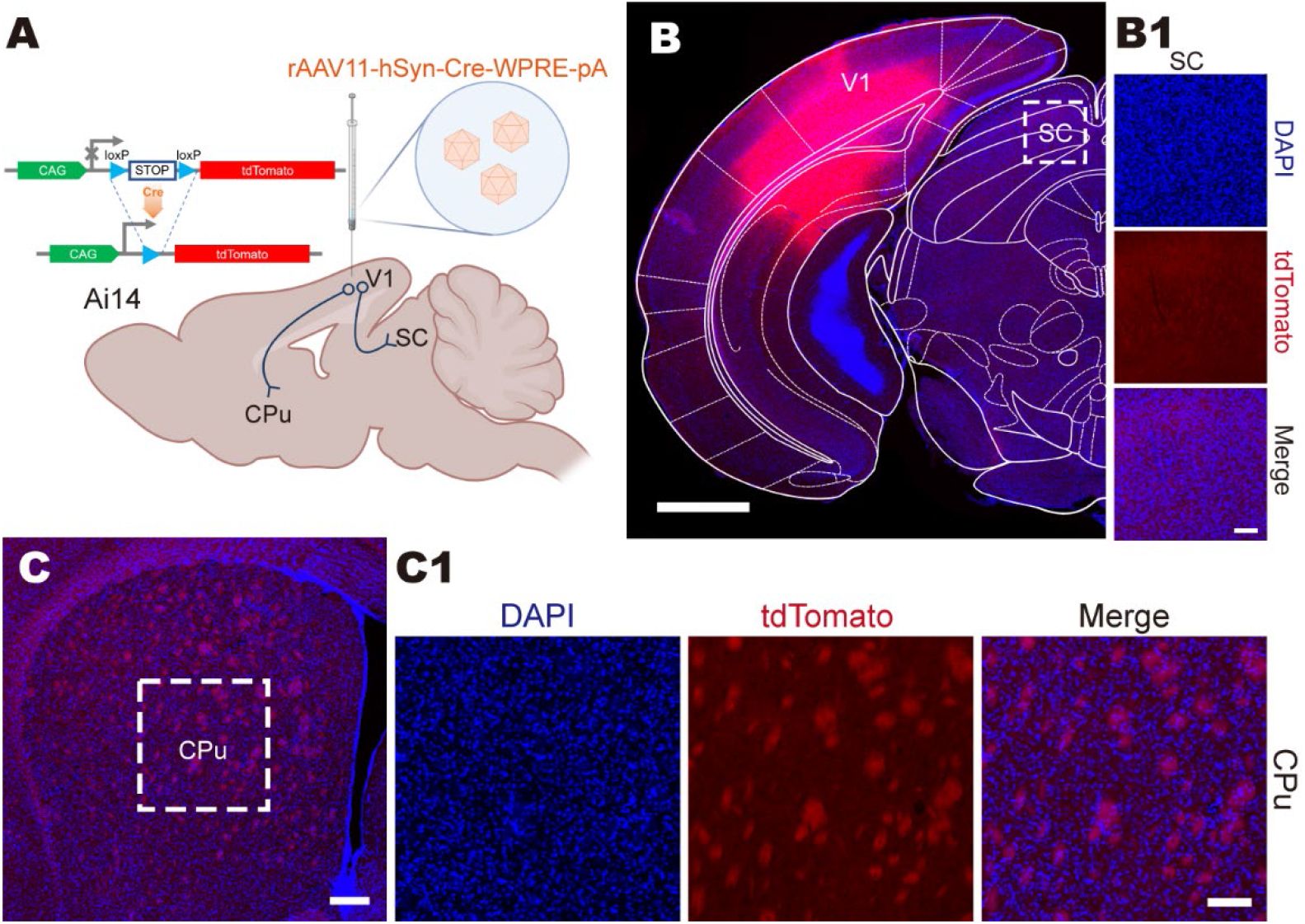
rAAV11 does not exhibit anterograde transsynaptic spread properties. (A) Connectivity diagram shows the injection site (primary visual cortex, V1) and connected upstream areas in superior colliculus (SC) and caudate-putamen (CPu), rAAV11-hSyn-Cre-WPRE-pA (2 × 10^9^ VG) was performed in Ai14 transgenic mice, in which the expression of tdTomato fluorescent reporter is Cre-dependent. (B) The tdTomato fluorescence expressed in injection site (V1). Scale bar = 1 mm. (B1) No tdTomato fluorescence signal is detected in superior colliculus (SC) area, which is the downstream brain area of V1. Scale bar = 1 mm. (C) No tdTomato fluorescence signal is detected in caudate-putamen (CPu) area, which is the downstream of V1. Scale bar = 200 μm for C and 100 μm for C1.

### rAAV11 can trace projection neurons that rAAV2-retro does not easily transduce

As an effective and practical retrograde viral tracer, rAAV2-retro is widely used in the analysis and manipulation of different types of neural circuitry [41, 42], as well as disease modeling and therapeutic evaluation [43]. To estimate the performance of rAAV11 retrograde tagging, we compared rAAV11 and rAAV2-retro by injecting an equal amount of rAAV2-retro and rAAV11 carrying different fluorescent reporter genes. rAAV11-EF1α-EGFP and rAAV2-retro-EF1α-mCherry are mixed in equal amounts and injected into CPu (Fig.3), ventral hippocampus (vHPC) (Fig.4) and periaqueductal gray (PAG) (Fig.5), respectively.

**Fig.3.**
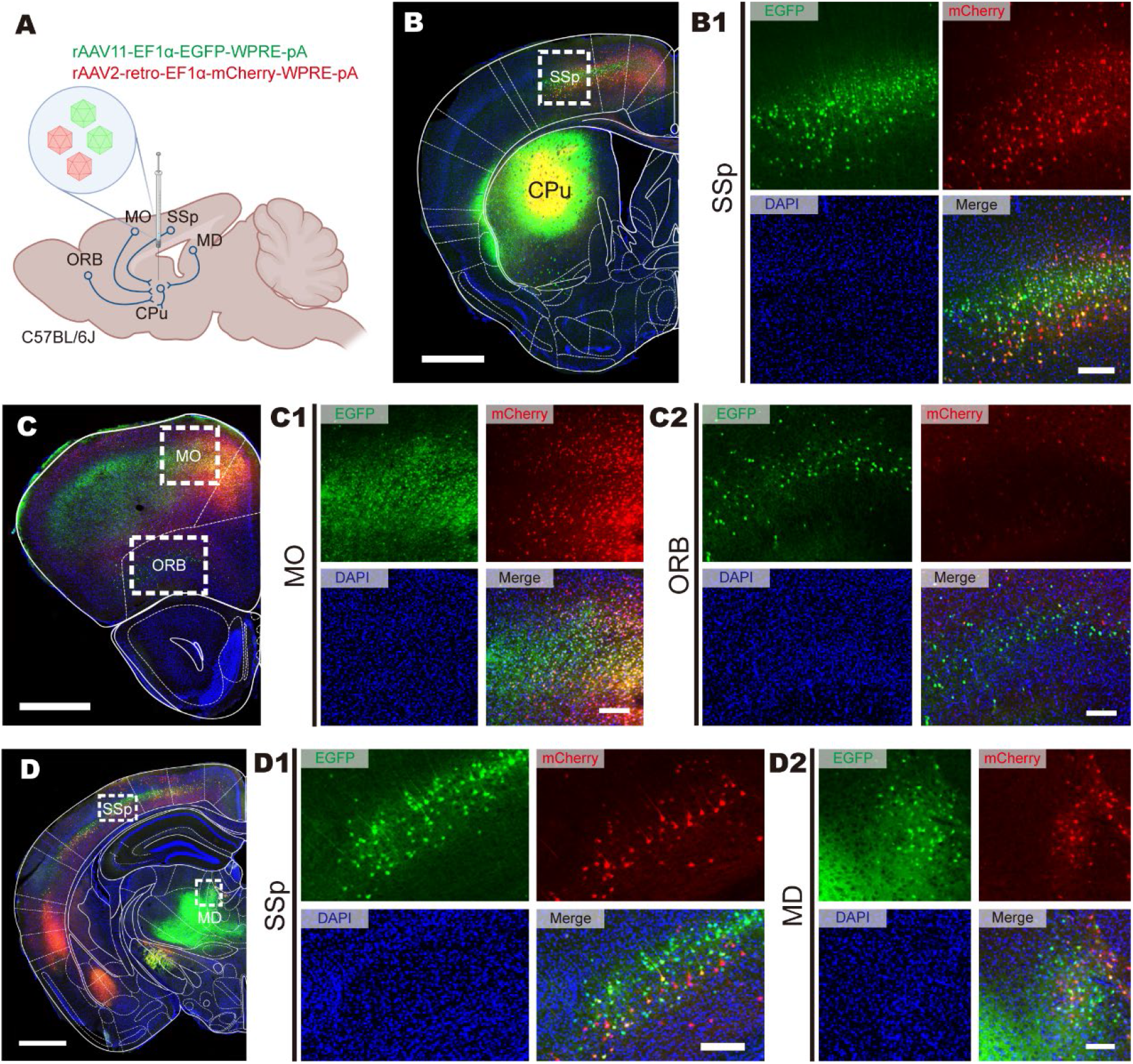
Retrograde infection tropism comparison of rAAV11 and rAAV2-retro injected into the CPu. (A) Connectivity diagram shows the injection site (caudate-putamen, CPu) and connected upstream areas in primary somatosensory area (SSp), somatomotor areas (MO), orbital area (ORB), and mediodorsal nucleus of thalamus (MD), rAAV11-EF1α-EGFP-WPRE-pA and rAAV2-retro-EF1α-mCherry-WPRE-pA (3 × 10^9^ VG in total) was performed in C57BL/6J mice. (B) Fluorescence distribution of EGFP (rAAV11) and mCherry (rAAV2-retro) expression in injection site CPu. Scale bar = 1 mm. (B1) The fluorescence distribution of EGFP and mCherry expression in SSp. Scale bar = 150 μm. (C) Fluorescence expression of neurons after virus infection in the anterior side of injection site. Scale bar = 1 mm. Representative images reveal that the retrograde labeling patterns of rAAV11-EF1α-EGFP and rAAV2-retro-EF1α-mCherry are quite different in many regions, such as the MO (C1, Scale bar = 150 μm) and ORB (C2, Scale bar = 150 μm). (D) Fluorescence expression of neurons after virus infection in the posterior side of injection site. Scale bar = 1 mm. Representative images reveal that the retrograde labeling patterns of rAAV11-EF1α-EGFP and rAAV2-retro-EF1α-mCherry are quite different in many regions, such as the SSp (D1, Scale bar = 150 μm) and MD (D2, Scale bar = 100 μm).

**Fig.4.**
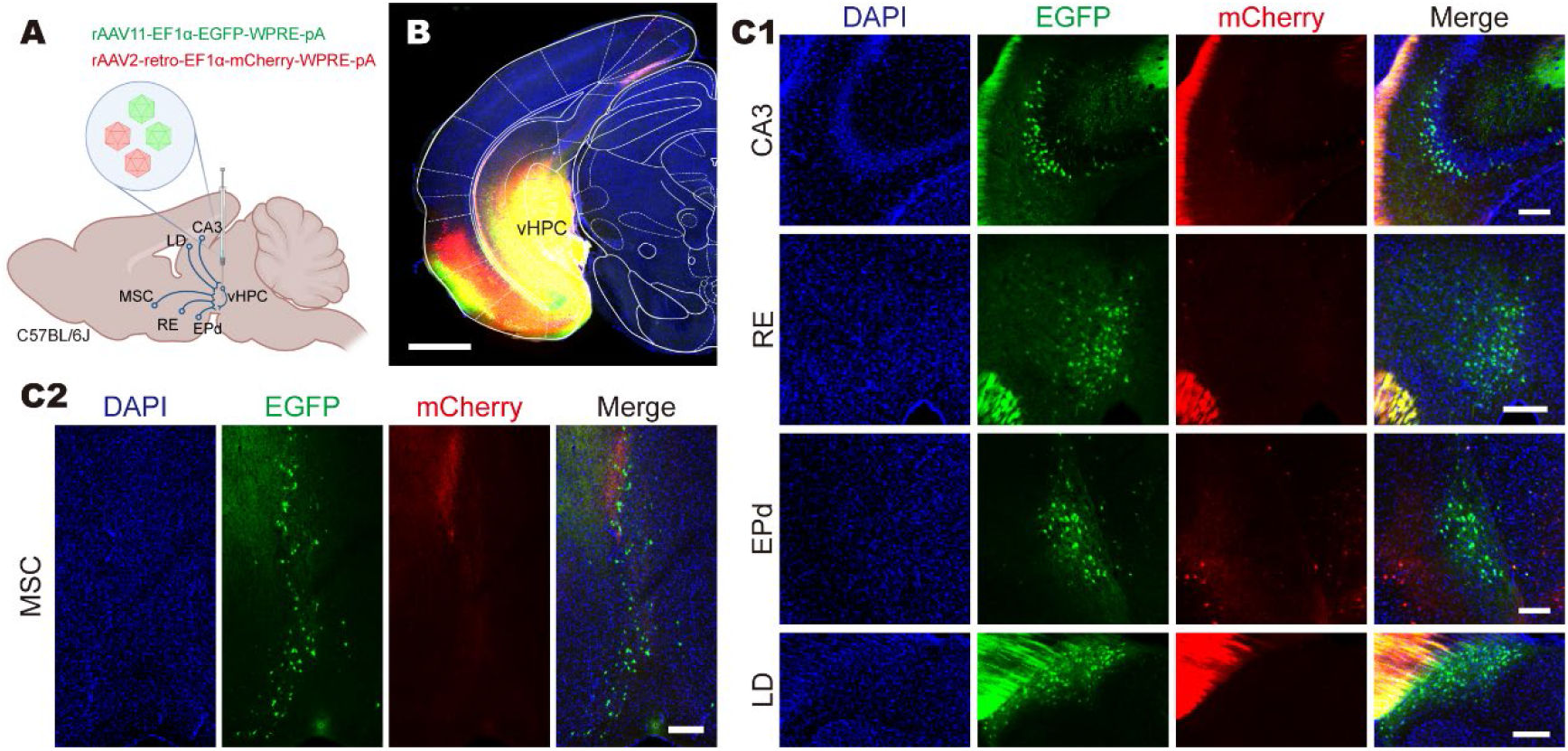
Retrograde infection tropism comparison of rAAV11 and rAAV2-retro injected into the vHPC. (A) Connectivity diagram shows the injection site (ventral hippocampus, vHPC) and connected upstream areas in hippocampal field CA3 (CA3), nucleus of reuniens (RE), endopiriform nucleus dorsal part (EPd), lateral dorsal nucleus of thalamus (LD), and medial septal complex (MSC), rAAV11-EF1α-EGFP and rAAV2-retro-EF1α-mCherry viruses were mixed at the particle ratio of 1:1 (3 × 10^9^ VG in total) and were injected into vHPC of C57BL/6J mice. (B) Fluorescence distribution of EGFP (rAAV11) and mCherry (rAAV2-retro) expression in injection site vHPC. Scale bar = 1 mm. (C) Representative images reveal that the retrograde labeling patterns of rAAV11-EF1α-EGFP and rAAV2-retro-EF1α-mCherry are quite different in many regions, such as the CA3, RE, EPd, LD (C1, Scale bar = 150 μm) and MSC (C2, Scale bar = 200 μm).

**Fig.5.**
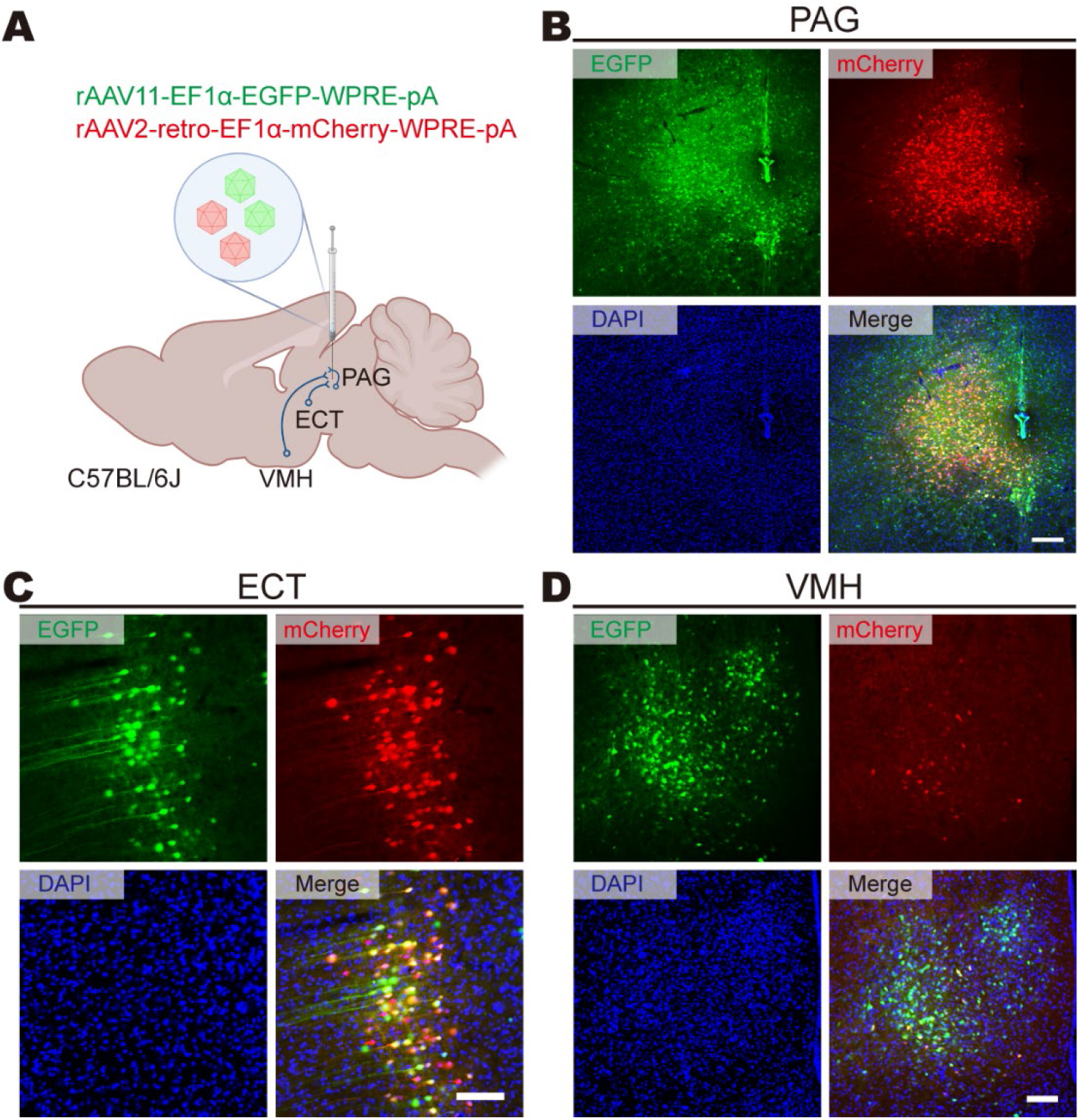
Retrograde infection tropism comparison of rAAV-Y and rAAV2-retro injected into the PAG. (A) Connectivity diagram shows the injection site (periaqueductal gray, PAG) and connected upstream areas in ectorhinal area (ECT) and ventromedial hypothalamic nucleus (VMH), rAAV11-EF1α-EGFP and rAAV2-retro-EF1α-mCherry viruses were mixed at the particle ratio of 1:1 (1.8 × 10^9^ VG in total) and were injected into PAG of C57BL/6J mice. (B) Fluorescence distribution of EGFP (rAAV11) and mCherry (rAAV2-retro) expression at injection site PAG. Scale bar = 200 μm. (C) EGFP (rAAV11) and mCherry (rAAV2-retro) expression in ECT. Scale bar = 100 μm. (D) EGFP (rAAV11) and mCherry (rAAV2-retro) expression in VMH. Scale bar = 100 μm.

At the CPu injection site, it can be observed that rAAV11 has a more extensive spreading range than rAAV2-retro (Fig.3 B), in the upstream SSp brain area of CPu, both tracers can be detected, but rAAV11 label more cells and tends to infect the fourth layer, while rAAV2-retro prefer to infect the fifth layer (Fig.3 B1 D1), similar infection tendency can also be observed in MO (Fig.3 C1) and MD (Fig.3 D2), still, rAAV11 has a more extensive infection range. In particular, in the orbital area (ORB), there were far more cells labeled with rAAV11 than rAAV2-retro, and rAAV2-retro only labeled a few cells. A similar pattern is illustrated in Figure 4, when injected in the vHPC area (Fig.4 A, B). rAAV11 has excellent retrograde tracing in hippocampal field CA3 (CA3), nucleus of reuniens (RE), endopiriform nucleus dorsal part (EPd), lateral dorsal nucleus of thalamus (LD), and medial septal complex (MSC), while rAAV2-retro label few cells (Fig.4 C1, C2). The periaqueductal gray (PAG) is a critical structure in the propagation and modulation of pain, sympathetic responses, as well as the learning and action of defensive and aversive behaviors [44]. We also tested the ability of rAAV11 and rAAV2-retro to retrogradely label the upstream brain regions of PAG (Fig.5). Although a comparable level of tracing effect can be seen in ectorhinal area (ECT) (Fig.5 C), rAAV11 still shows a better tracing effect in ventromedial hypothalamic nucleus (VMH) and offers a more extensive spread range in PAG.

### Use rAAV11 to trace the upstream of the SSp area combined with Ai14 transgenic mice

Humans and other primates use their hands to engage in tactile discrimination that directs choices and actions. The vast majority of such decisions are based on stimuli which are presented to the sites by hand or hands, the transformation of these information sources dispersed from receptors in distinct plates of the skin into a percept presupposes integration into the central nervous system, SSp played an essential role in this process [45]. As a proof of principle, with Ai14 transgenic mice, we use rAAV11-Cre to trace the upstream projection neurons of SSp (Fig.6 A, B). The tdTomato signal is visible in ventral anterior-lateral complex of the thalamus (VAL) and ectorhinal area (ECT) (Fig.6 C).

**Fig.6.**
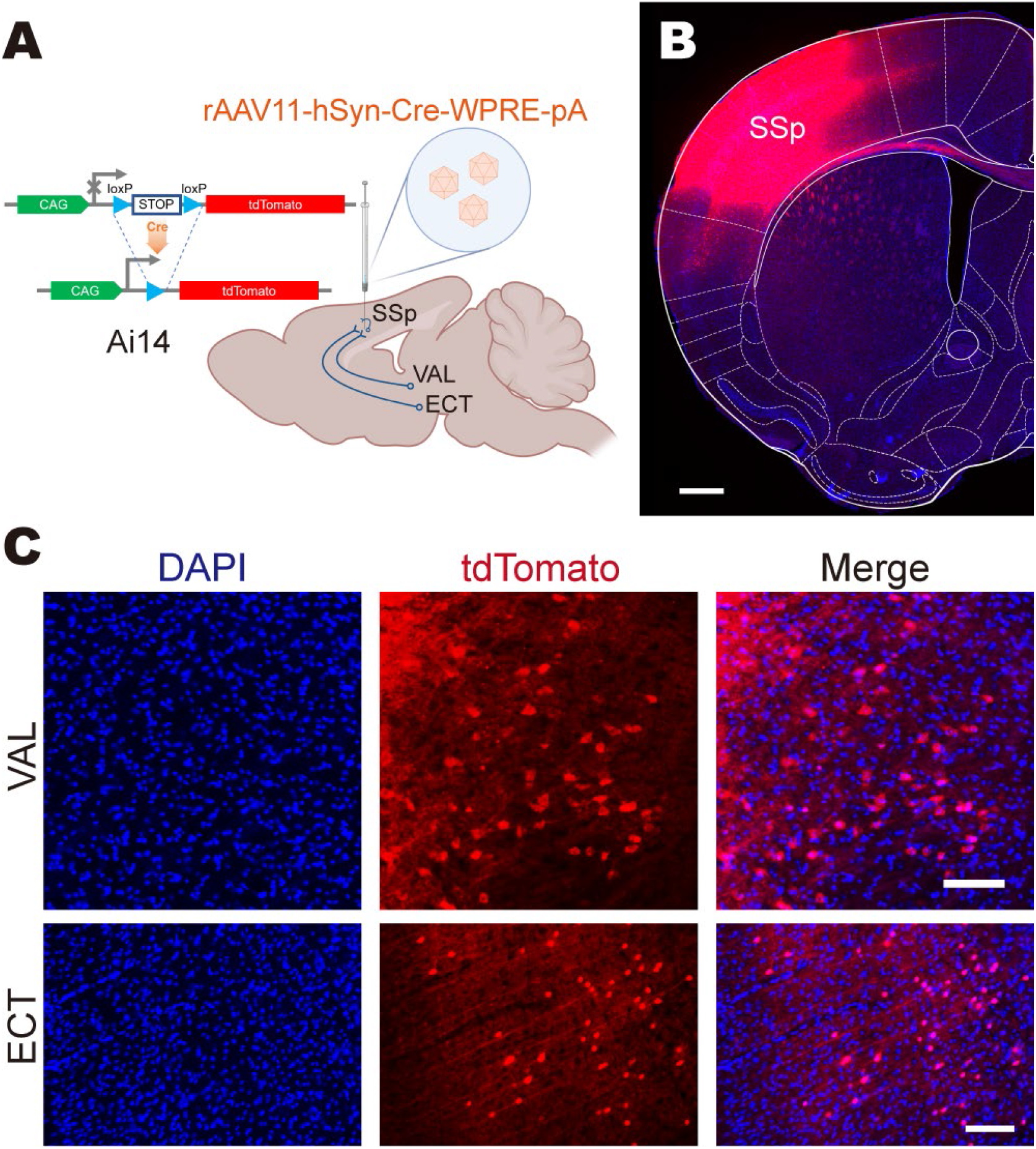
Use the rAAV11 to trace the upstream input network of the SSp. (A) Connectivity diagram shows the injection site (primary somatosensory area, SSp) and connected upstream areas in ventral anterior-lateral complex of the thalamus (VAL) and ectorhinal area (ECT), rAAV11-hSyn-Cre (2 × 10^9^ VG) was performed in Ai14 transgenic mice, in which the expression of tdTomato fluorescent reporter is Cre-dependent. (B) The tdTomato fluorescence expressed in injection site (SSp). Scale bar = 500 μm. (C) The tdTomato fluorescence expressed in upstream areas such as VAL and ECT. Scale bar = 100 μm.

### Using rAAV11 for functional circuit interrogation

GCaMP is a genetically encoded calcium sensor consisting of a circular GFP, calmodulin (CaM), and a peptide chain (M13), which in its natural conformation shows only poor fluorescence. In the presence of calcium, CaM undergoes a structural change that entails a rapid increase in fluorescence [46]. GCaMP6m reliably detected single action potentials in neuronal somata and orientation-tuned synaptic calcium transients in individual dendritic spines [47]. The dopamine projection from the ventral tegmental area (VTA) to nucleus accumbens (NAc) is critical for motivation to work for rewards and reward-driven learning [48]. To verify the application of rAAV11 in functional circuit interrogation, the experiment was conducted using a retrogradely induced GCaMP6m expression, using an injection of an rAAV11-Cre vector in the NAc combined with an infusion of a Cre-inducible GCaMP6m vector in the VTA (Fig. 7A) of C57BL/6J mice. We set up sucrose solution as a reward in the behavior box of the mice (Fig.7 A, B), when mice licked sugar water to get rewards, VTA neurons projecting to the NAc were activated (Fig.7 C), then the calcium ion signal could be detected (Fig.7 D, E).

**Fig.7.**
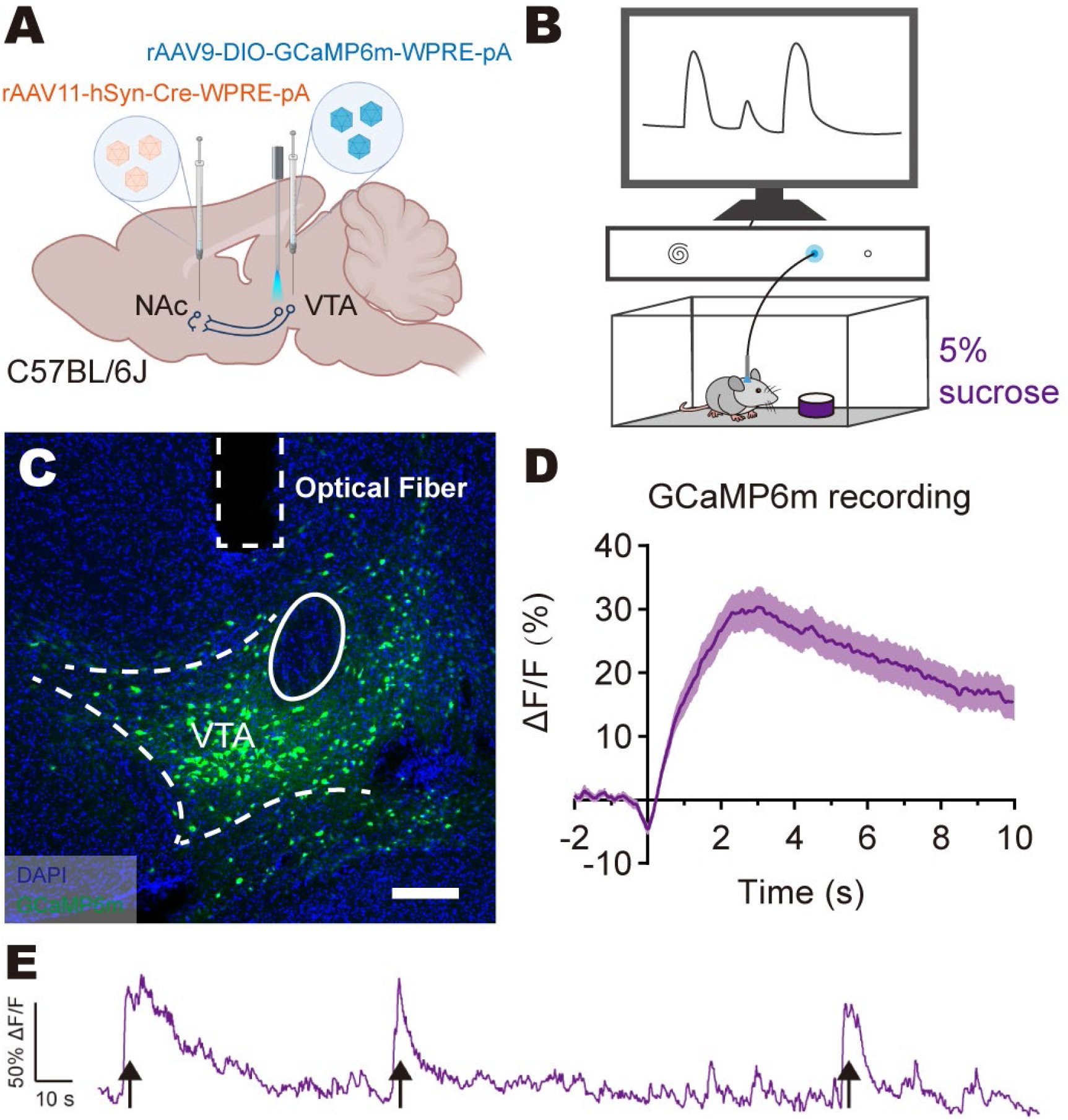
rAAV11 can be used for analyzing functional network. (A) Experimental injection paradigm for functional verification of the afferents from the ventral tegmental area (VTA) to the nucleus accumbens (NAc), rAAV11-hSyn-Cre (2 × 10^9^ VG) was injected in NAc and rAAV9-DIO-GCaMP6m (2 × 10^9^ VG) in VTA, optical fiber for calcium ion signal detecting was implanted in VTA. (B) The experimental set up. (C) The GCaMP6m fluorescence expressed in VTA. Scale bar = 200 μm. (D) Behavior-triggered average of ΔF/F signals. (E) A representative trace of GCaMP6m ΔF/F signal with black arrows indicating when the animal licked the sucrose solution.

### rAAV11 effectively target astrocytes

Glial cells are critical responders to central nervous system injury and equal the number of neurons in the adult human brain [49, 50]. Furthermore, astrocytes are the most abundant glial cell type in the CNS and they play a crucial role in the pathogenesis of spinal cord injury [51]. However, some serotypes of AAV cannot target glial cells, the viral tracers for specifically targeting astrocytes are still lacking. We tested the targeting ability of rAAV11 to astrocytes in dentate gyrus (DG) (Fig.8 A, B). By carrying the EGFP reporter gene initiated by GfaABC1D promoter, we observed that the green fluorescent protein and astrocytes have a reasonable co-labeling (Fig.8 C).

**Fig.8.**
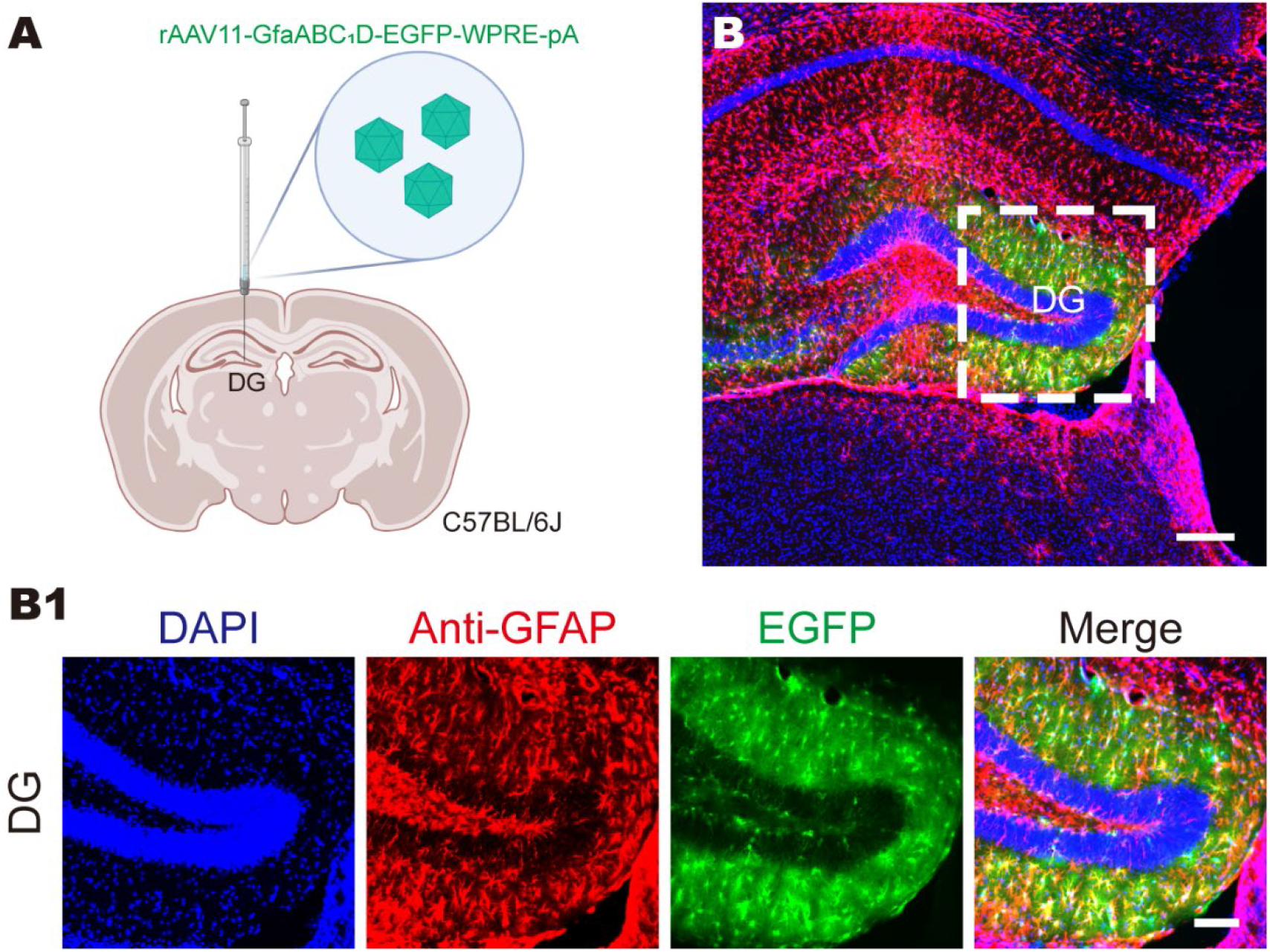
rAAV11 effectively target astrocytes. (A) Schematic diagram of virus injection, rAAV11-GfaABC1D-EGFP (2 × 10^9^ VG) virus was injected into dentate gyrus (DG) area of C57BL/6J mice. (B) EGFP expression in injection site DG with anti-GFAP indicating astrocytes. Scale bars = 200 μm for B and 100 μm for B1.

## Discussion

Currently, AAV is among the most commonly used viruses for neuroscience research [52] and the leading platform for *in vivo* delivery of gene therapies [53], have been extensively used as vehicles for gene transfer to the nervous system enabling gene expression and knockdown, gene editing [54, 55], circuit modulation [56, 57], *in vivo* imaging [58, 59], disease model development [60], and the evaluation of therapeutic candidates for the treatment of neurological diseases [18]. Focus on the application of AAV vectors in central nervous system related issues, in addition to the delivery of the gene of interest by *in situ* injection, the infection types of AAV mainly include across the blood-brain barrier via intravenous injection [61–64], anterograde transsynaptic spread [37], and retrograde labeling by axon terminal absorption [20–23, 65].

There are various natural serotypes of AAV, which belong to different pedigrees and have their unique tropism [25–27, 66], this is one of the reasons why AAV is widely used. Our research has tested the rarely used natural serotype AAV11, which capsid has not been artificially modified, and found that it has a highly efficient retrograde effect. AAV2-retro was promoted and applied, and the AAV9-retro [21], AAV2-MNM004 [22], and AAV9-SLR [23] were obtained by engineering modification later, but the effect of retrograde can only be equal to or inferior to AAV2-retro. Therefore, there are two points worth noting in our research: first, the retrograde capability of rAAV11 has been better than rAAV2-retro to a certain extent (Fig.3, Fig.4, Fig.5); second, as a natural serotype, the capsid of AAV11 still has excellent potential for further technical upgrades. In addition, we confirmed that rAAV11 does not have anterograde transsynaptic properties (Fig.2), and can carry functional elements for genetic manipulation (Fig.6, Fig.7).

Especially, we found that rAAV11 can efficiently target astrocytes (Fig.8). Although astrocytes have traditionally been described as playing a supportive role for neurons, they have recently been recognized as active participants in the development and plasticity of dendritic spines and synapses [67]. Astrocyte dysfunction has in recent years proven to be a common crossroads in neurodegenerative disorders such as stroke [68], Alzheimer’s disease [69], Parkinson’s disease [70] and Huntington’s disease [71], alteration in astrocytic glutamate uptake is a core feature of multiple neurodegenerative disorders [67, 72], with the in-depth research on astrocytes, rAAV11 may be used as a powerful candidate virus tool for scientific research or gene therapy.

Based on the above nervous system infection tropism, rAAV11 can become a new engineering target for virus vector designers. AAV capsid engineering offers great potential and the successful promotion of mutation vectors such as AAV2-retro benefits from this. There are at least four main approaches currently used towards this: directed evolution [20, 22, 61, 62, 73, 74], rational design [75–79], *in silico* evolutionary lineage analysis [80, 81], and chemical conjugation [82–88]. The “natural occurring” of AAV11 capsid has become one of its advantages, with reference to the previous experience of AAV capsid engineering, it might be possible to obtain a better AAV11 mutant version through artificial upgrades, whether in retrograde labeling or astrocyte targeting.

## Material and Methods

### Plasmids construction and virus manufacturing

To obtain the pAAV2/11 plasmid, AAV11 *Cap* sequence retrieved on Genbank (accession number: AY631966.1) was synthesized as a template for amplifying AAV11 *Cap* fragment using PrimeSTAR HS DNA Polymerase (Takara, R040A), After being digested by using SwaI and AgeI (New England Biolabs) restriction endonucleases, the AAV11 *Cap* fragment was inserted into pAAV-RC2/1 vector (Addgene, #112862) by T4 DNA ligase (New England Biolabs, M0202M), then the ligation product was transformed into Stbl3 chemically competent *E. coli*, the positive clone was picked after PCR identification and the plasmid was extracted to obtain pAAV2/11.

The plasmids carrying the foreign gene of interest (e.g., pAAV-hSyn-Cre-WPRE-pA, pAAV-EF1α-EGFP-WPRE-pA, pAAV-EF1α-mCherry-WPRE-pA, pAAV-EF1α-DIO-GCaMP6m-WPRE-pA or pAAV-GfaABC_1_D-EGFP-WPRE-pA) with pAAV-RC2/11 (or pAAV-RC2-retro or pAAV-RC9) and pAdDeltaF6 (Addgene plasmid #112867) were co-transfected into HEK-293T cells at the molecular ratio of 1:1:1, samples were harvested at 72 hours post-transfection and purified by iodixanol gradient ultracentrifugation [29, 89, 90]. The purified rAAVs were titred by qPCR using the iQ SYBR Green Supermix kit (Bio-Rad, 1708884) and diluted to 1.0 × 10^13^ VG/mL. All viral vectors were aliquoted and stored at −80 ℃ until use.

### Research animals

Adult male (8–10 weeks old) C57BL/6J mice (Hunan SJA Laboratory Animal Company) and Ai14 transgenic mice (The Jackson Laboratory) were used for experiments. The mice were housed in the appropriate environment with a 12/12-h light/dark cycle, water and food were supplied *ad libitum*. All the surgical and experimental procedures were performed following the guidelines formulated by the Animal Care and Use Committee of Innovation Academy for Precision Measurement Science and Technology, Chinese Academy of Sciences.

### Stereotaxic AAV injection

Briefly, the mice were deeply anaesthetized using 1% pentobarbital intraperitoneally (i.p., 50 mg/kg body weight). The stereotactic injection coordinates were selected according to Paxinos and Franklin’s *The Mouse Brain in Stereotaxic Coordinates*, 4th edition [91], animals were placed on a stereotactic frame (RWD, 68030). A small volume of the virus was injected into the CPu (anterior-posterior-axis (AP) +0.75 mm from bregma, medial-lateral-axis (ML) ±2.00 mm, and dorsal-ventral-axis (DV) −2.00 mm), V1 (AP −3.90 mm from bregma, ML ±2.6 mm, and DV −1.30 mm), vHPC (AP − 3.16 mm from bregma, ML ±2.95 mm, and DV −4.10 mm), PAG (AP −4.00 mm from bregma, ML ±0.26 mm, and DV −2.60 mm), SSp (AP +0.50 mm from bregma, ML ±3.00 mm and DV −2.00 mm), VTA (AP −3.20 mm from bregma, ML ±0.45 mm, and DV −4.30 mm), NAc (AP +1.50 mm from bregma, ML ±1.10 mm, and DV −4.60 mm), and DG (AP −2.15 mm from bregma, ML ±1.30 mm, and DV −2.00 mm), respectively, using a pulled glass capillary with stereotaxic injector (Stoelting, 53311) at a slow rate of 0.03 μL/min. After the injection was completed, the capillary was left for an additional 10 minutes before slowly being withdrawn completely. After surgery, animals were allowed to recover from anaesthesia under a heating pad. At three weeks post-injection, the mice were sacrificed. Mice were deeply anesthetized with an overdose of 5% chloral hydrate and transcardially perfused ice-cold PBS (pH 7.4) followed by 4% paraformaldehyde. After overnight post fix in 4% paraformaldehyde solution, brains were dehydrated in 30% sucrose solution for one day.

### Slice preparation, immunohistochemistry, and imaging

Slice preparation and imaging were completed according to the previously reported methods [21]. Coronal sections (40 μm) were cut on a microtome (Thermo Fisher Scientific), collected in anti-freeze fluid, and stored at −20 °C for further use. For staining the GFAP, the primary antibody used were goat anti-GFAP (1:800, Abcam), amplified with secondary antibody rabbit anti-goat IgG conjugated with Cy3 (1:400, Jackson), and fixed slices stained with DAPI (1:4000), then washed with PBS three times, followed by sealing with 70% glycerol. Imaging was performed using the Olympus VS120 Slide Scanner microscope (Olympus).

### Imaging of neuronal population activity in vivo following delivery of GCaMP6m

rAAV11-hSyn-Cre (2 × 10^9^ VG) and rAAV9-DIO-GCaMP6m (2 × 10^9^ VG) was injected into the NAc and VTA, respectively, and optical fiber (core diameter: 200 μm, numerical aperture: 0.37, Inper, China) was implanted in VTA. Mice had visually identifiable GCaMP6m-expressing cells in VTA two weeks post-injection. Before recording, the mice were handled for 3–5 min at least 3 days and then were habituated to the fiber patch cord and a chamber (20×20×22 cm) for 10 min. All mice were water deprived for 24 h before placed in the chamber, which was equipped with a cup filled with 5 % (w/v) sucrose solution. Then the mice were tested when searching for sucrose rewards. Calcium transients during behavior were recorded by exciting GCaMP6m at 470 nm using the fiber photometry system (ThinkerTech, Nanjing, China). Data were analysed using MATLAB (MathWorks) code and Prism (GraphPad software).

## Acknowledgments

We are grateful to Liting Luo and Lingling Xu (Core Facility Center, Innovation Academy for Precision Measurement Science and Technology, Chinese Academy of Sciences) for technical assistance with centrifugation and microscope imaging. Schematic diagrams in the pictures of this article were created with BioRender.com. This work was supported by the National Natural Science Foundation of China (31830035, 31771156, 21921004), the Key-Area Research and Development Program of Guangdong Province (2018B030331001), the Strategic Priority Research Program of the Chinese Academy of Sciences (XDB32030200) and the Shenzhen Key Laboratory of Viral Vectors for Biomedicine (ZDSYS20200811142401005).

## Authors’ contributions

KL and FX contributed to the study idea and design; FX, CY and JW contributed to funding acquisition and resources; ZH, NL, JK, LL, WM, SP, ZX, WZ and YQ performed the experiments and data acquisition; ZH, NL, KL and YW accomplished data analysis; ZH, KL and FX drafted the manuscript, and contributed to review and editing. All authors read and approved the final manuscript.

